# Durability of the Neutralizing Antibody Response to mRNA Booster Vaccination Against SARS-CoV-2 BA.2.12.1 and BA.4/5 Variants

**DOI:** 10.1101/2022.07.21.501010

**Authors:** Panke Qu, Julia N. Faraone, John P. Evans, Yi-Min Zheng, Claire Carlin, Gerard Lozanski, Linda J. Saif, Eugene M. Oltz, Richard J. Gumina, Shan-Lu Liu

## Abstract

The recent emergence of the SARS-CoV-2 BA.4/5 and BA.2.12.1 variants has led to rising COVID-19 case numbers and concerns over the continued efficacy of mRNA booster vaccination. Here we examine the durability of neutralizing antibody (nAb) responses against these SARS-CoV-2 Omicron subvariants in a cohort of health care workers 1-40 weeks after mRNA booster dose administration. Neutralizing antibody titers fell by ∼1.5-fold 4-6 months and by ∼2.5-fold 7-9 months after booster dose, with average nAb titers falling by 11-15% every 30 days, far more stable than two dose induced immunity. Notably, nAb titers from booster recipients against SARS-CoV-2 BA.1, BA.2.12.1, and BA.4/5 variants were ∼4.7-, 7.6-, and 13.4-fold lower than against the ancestral D614G spike. However, the rate of waning of booster dose immunity was comparable across variants. Importantly, individuals reporting prior infection with SARS-CoV-2 exhibited significantly higher nAb titers compared to those without breakthrough infection. Collectively, these results highlight the broad and stable neutralizing antibody response induced by mRNA booster dose administration, implicating a significant role of virus evolution to evade nAb specificity, versus waning humoral immunity, in increasing rates of breakthrough infection.

## Introduction

The coronavirus disease 2019 (COVID-19) pandemic has had devastating impacts across the globe, with over 500 million confirmed cases and 6 million deaths worldwide since its emergence (World Health Organization, 2022). Vaccines against the causative agent, severe acute respiratory syndrome coronavirus 2 (SARS-CoV-2), were rapidly developed, including two mRNA vaccines, Moderna mRNA-1273 and Pfizer/BioNTech BNT162b2. mRNA vaccination has led to a reduction in the number of COVID-19 cases, as well as hospitalizations or deaths (Andrews et al., 2022; Chenchula et al., 2022; Scobie et al., 2021). The standard regimen for these vaccines includes two doses separated by about four weeks. This was later supplemented with an additional booster dose at least six months after the second dose. Recent studies have demonstrated that humoral immunity induced by two doses of mRNA vaccine wanes significantly over time, while a booster dose can compensate for these effects, albeit to a lesser extent against Omicron as compared to the Delta variant (Evans et al., 2022a; Qu et al., 2022; Richterman et al., 2022). Critically, the durability of immunity stimulated by booster vaccination is currently not well understood.

Over the course of the pandemic, SARS-CoV-2 has evolved to become more transmissible and less sensitive to humoral immunity, resulting in several major SARS-CoV-2 variants of concerns. Mutations such as K417N and E484K/A, present in the Beta, Gamma, and Omicron variants, have led to increased resistance to neutralizing antibodies (nAbs) (Ghimire et al., 2022; Rajpal et al., 2022). The Omicron variant in particular has exhibited the most substantial immune evasion due to the alarming number of amino acid mutations in its spike gene, totaling more than 30, with 16 concentrated in the receptor binding domain (RBD) (O’Toole et al., 2021). Several subvariants of Omicron have emerged, especially BA.4/5 and BA.2.12.1, creating new waves of COVID-19 across the globe. It has been established that these Omicron subvariants exhibit strong resistance to nAbs induced by two-dose mRNA vaccination, but this resistance can be partially overcome by booster vaccination (Evans et al., 2022b, 2022a; Qu et al., 2022; Richterman et al., 2022). There have been reports that protection provided by a booster dose can last at least 4 months post-vaccination (Ferdinands et al., 2022; Richterman et al., 2022), but the durability of booster induced immunity past this timepoint remains unclear. Additionally, the durability of booster protection against more recent Omicron subvariants BA.2.12.1 and BA.4/5 has yet to be investigated. To address this, we examine the nAb response in mRNA vaccinated and boosted health care workers (HCWs) against major circulating SARS-CoV-2 Omicron subvariants from 1 to 9 months post-booster administration. We observe modest waning of booster induced immunity that is dependent on prior COVID-19 status, while the Omicron sub variants, especially BA.4/5, exhibit strong neutralization resistance.

## Results

### Omicron subvariants BA.4/5 and BA.2.12.1 substantially evade booster-induced immunity

Given ongoing concern for the durability of protection offered by the mRNA vaccine booster doses, especially against Omicron variants, we examined neutralizing antibody (nAb) titers against SARS-CoV-2 in a longitudinal cohort of HCWs from The Ohio State University Wexner Medical Center in Columbus, Ohio. These HCWs provided serum samples every 3 months following administration of the second mRNA vaccine dose. HCWs received homologous vaccine and booster courses consisting of the Moderna mRNA-1273 (n = 24, Table S1) or Pfizer/BioNTech BNT162b2 (n = 22) mRNA vaccines. Due to variability in the timing of booster dose administration, we classified the HCW samples into 3 groups, i.e., 1-3 months post booster dose, 4-6 months post booster dose, and 7-9 post booster dose (**Table S1)**.

To examine the nAb responses against major circulating SARS-CoV-2 Omicron subvariants, we utilized our previously reported pseudotyped lentivirus neutralization assay (Zeng et al., 2020). We generated virus pseudotyped with spike protein from the ancestral D614G SARS-CoV-2 or major SARS-CoV-2 Omicron subvariants, including the original BA.1 Omicron variant, the recently dominant BA.2.12.1 variant, and the BA.5 variant currently rising in cases numbers in the United States (Centers for Disease Control and Prevention, 2022).

All Omicron subvariants exhibited a significant reduction in nAb titers, presented as 50% neutralization titers (NT_50_), relative to D614G at all timepoints tested (**Fig 1**). At 1–3-months post-booster, nAb titers against BA.1 were 4.7-fold (p<0.0001), BA.2.12.1 were 7.6-fold (p<0.0001), and BA.4/5 were 13.4-fold (p<0.0001) lower than D614G (**Fig 1A and 1D**). At 4-6 months post-booster, nAb titers against BA.1 were 5.6-fold (p<0.0001), BA.2.12.1 were 9.5-fold (p<0.0001), and BA.4/5 were 17.3-fold (p<0.0001) lower than D614G (**Fig 1B and 1E**). Finally, at the 7-9 months timepoint, nAb titers against BA.1 were 4.6-fold (p<0.0001), BA.2.12.1 were 7.0-fold (p<0.0001), and BA.4/5 were 13.4-fold (p<0.0001) lower than D614G (**Fig 1C and 1F**). Across all timepoints, BA.2.12.1 and BA.4/5 exhibited apparently reduced nAb titers compared to BA.1. For example, at the 1-3 month timepoint, nAb titers against BA.2.12.1 were 1.6-fold lower than against BA.1 (p=0.28) while for BA.4/5, nAb titers were 2.9-fold lower (p<0.0001) compared to BA.1 (**Fig 1A**). Similar fold differences were maintained throughout the later timepoints (4-6 months and 7-9 months post-booster) (**Fig 1B, 1E, 1C and 1F**). These results were consistent with several recent studies using samples collected from one single time point. No significant differences were observed for nAb titers between HCWs that received Pfizer (n=22) or Moderna (n=24) (p > 0.05), and Male (n=28) or Female HCWs (n=18) (p > 0.05) (**Fig S2A and S2B**).

**Figure 1:**
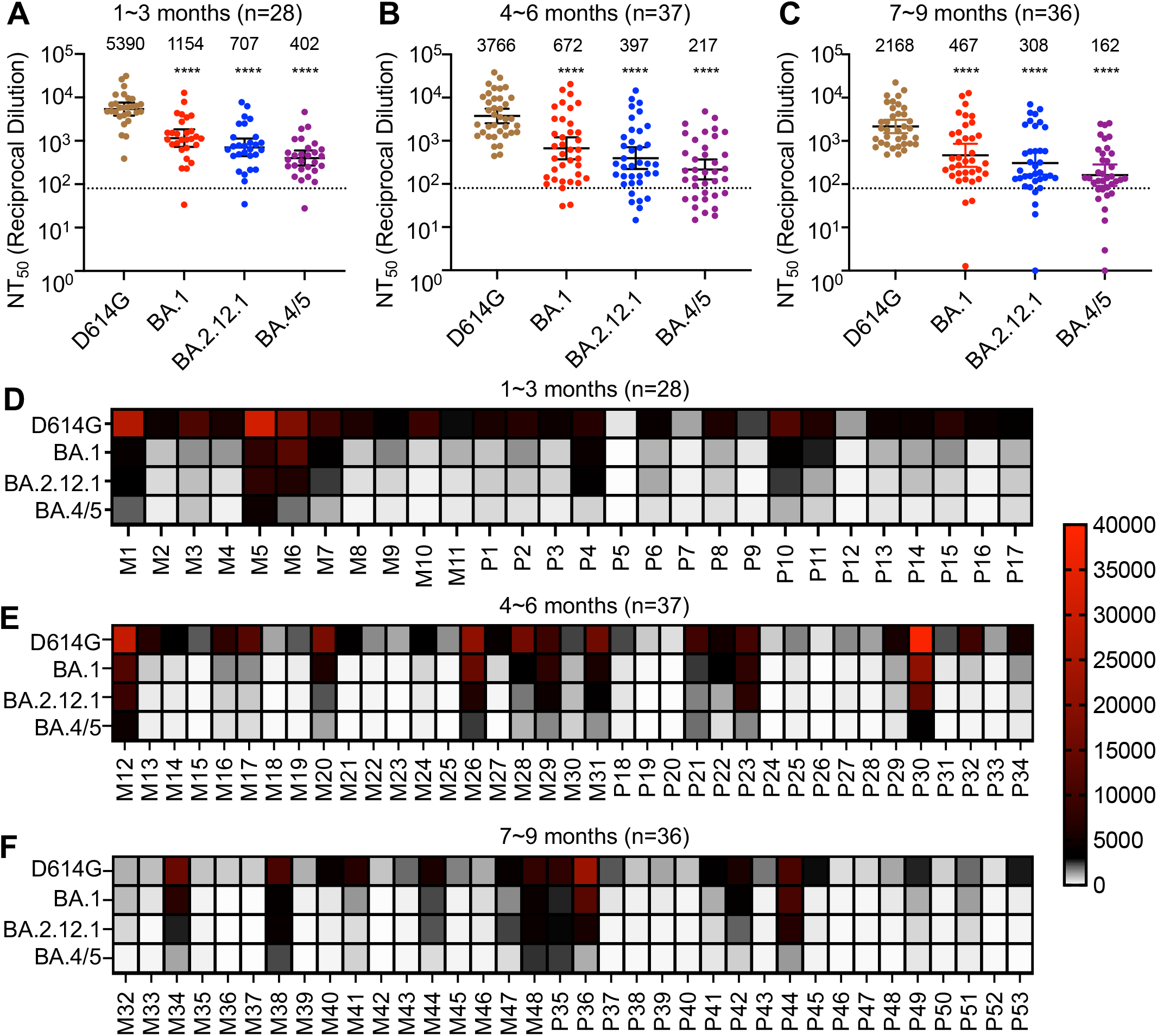
Omicron subvariants BA.4/5 and BA.2.12.1 exhibit strong escape from neutralization that is maintained over time post booster vaccination. These plots depict the nAb titers for the ancestral SARS-CoV-2 spike with the D614G mutation and the Omicron subvariants BA.1, BA.2.12.1, and BA.4/5 in serum samples from HCWs collected at **(A)** 1-3 months, **(B)** 4-6 months, and **(C)** 7-9 months after receiving a booster dose of mRNA vaccine. Dots represent individual samples while the horizontal dashed line represents the limit of detection. Geometric means of the NT_50_ values are provided at the top of the graph for each of the variants within each timepoint. Error bars represent 95% confidence intervals. **(D-F)** Corresponding heatmaps depicting NT_50_ values for each individual receiving either Moderna (M) or Pfizer (P) mRNA vaccine against each variant sorted by timepoint post-booster, **(D)** 1-3 months, **(E)** 4-6 months, and **(F)** 7-9 months. Significance values in **(A-C)** represent comparisons to D614G calculated with one-way repeated measures ANOVA with Bonferroni’s multiple testing correction. P-values are indicated as ****p<0.0001.

### Durability of mRNA booster vaccination decays over time

To determine the durability of the mRNA booster over the time course, we analyzed nAb titers post booster dose administration for each of the variants. As would be expected, the strength of virus neutralization against all 4 variants decreased over time, with 2.3-2.5-fold drop from 1-3 month to 7-9 month post booster vaccination. Correlative analyses showed an average of 15.3% (p = 0.0003), 13.5% (p = 0.037), 11.1% (p = 0.092), and 12.3% (p = 0.037) decline in nAb titers per 30 days for the D614G, BA.1, BA.2.12.1, and BA.4/5 variants, respectively (**Fig 2A-2D**). The similar rate of decay seems to indicate that the waning of neutralizing antibody responses following booster vaccination is not variant dependent.

**Figure 2:**
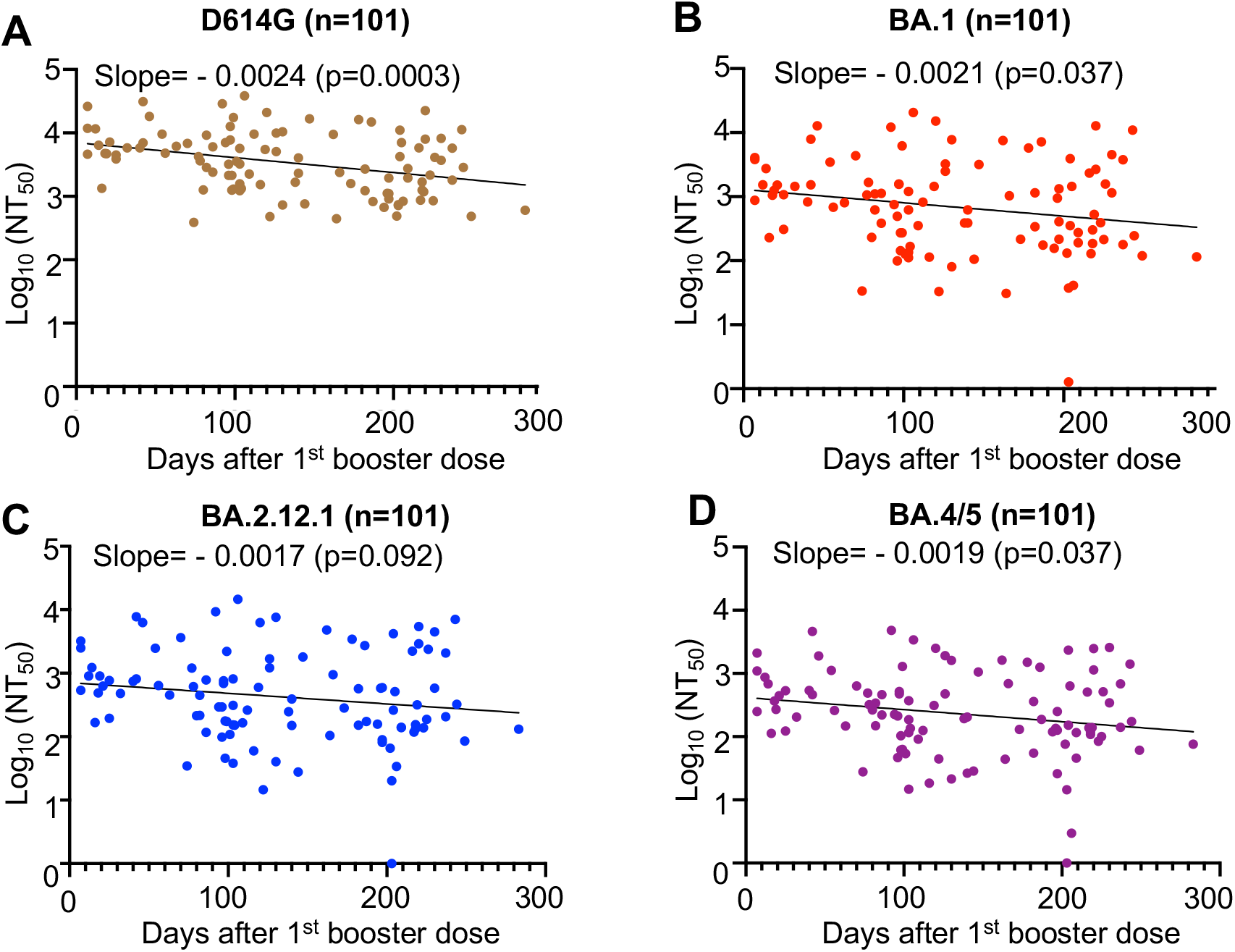
Durability of booster-induced nAb titers wanes over time. nAb titers against D614G and the Omicron subvariants BA.1, BA.2.12.1, and BA.4/5 in serum from HCWs collected after receiving a booster dose of mRNA vaccine are depicted as a function of number of days after receiving the booster. Each dot represents an individual sample, the line represents the best linear fit for the trend of nAb titer over time. Significance and slope for the trendline are listed at the top of each graph. Correlative analysis of nAb titers and time post booster dose administration was made using a least-squares fit linear regression model. Exact p-values are noted.

### Breakthrough infection enhances the durability of immunity

We next examined the impact of breakthrough infection on the durability of booster vaccine-induced immunity. Over the course of the study, 14 HCWs experienced breakthrough infections including 9 HCWs infected during the Omicron waves. Overall, nAb titers were 2-6-fold higher for HCWs that experienced breakthrough infections at the 1-3 month (p < 0.05), 4-6 month (p < 0.0001), and 7-9 month (p < 0.0001) post-booster ranges (**Fig 3A**). In particular, COVID-19 positive HCWs exhibited enhanced nAb titers against the Omicron subvariants at both the 4-6 and 7-9 month timepoints (**Fig 3A**). This indicates that breakthrough infection can enhance both nAb titers and the breadth of the nAb response.

**Figure 3:**
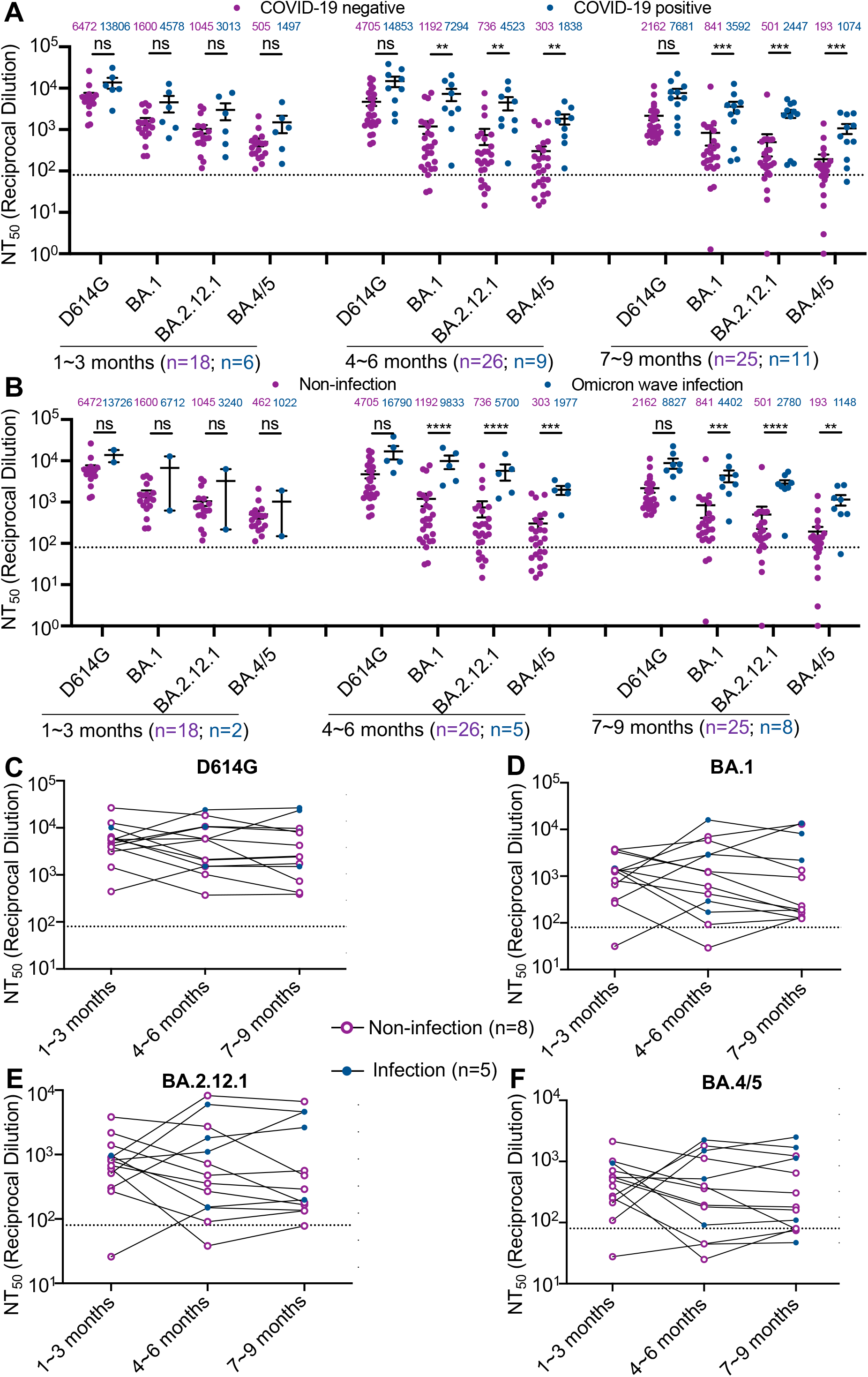
Breakthrough infection with SARS-CoV-2 increases and maintains nAb titers against Omicron subvariants. Depicted in this plot are the nAb titers against D614G and the Omicron subvariants BA.1, BA.2.12.1, and BA.4/5 in sera from HCWs collected after receiving a booster dose of mRNA vaccine separated by self-reported COVID-19 infection status. **(A)** Neutralizing antibody titers are displayed for HCWs that were previously diagnosed with COVID-19 versus those that remained uninfected throughout the study, sorted by the timepoints 1-3, 4-6, and 7-9 months-post booster dose. **(B)** Neutralizing antibody titers are displayed for HCWs that were infected during the Omicron wave in Columbus, Ohio versus those that remained uninfected throughout the study at the same timepoints. **(C-F)** Neutralizing antibody titers are displayed for 13 HCWs which provided all 3 sample collection timepoints. The NT_50_ values for each individual over time are depicted for each of the variants. Here, purple, open circles represent samples collected prior to any COVID-19 diagnosis while blue, closed circles represent samples that were collected after COVID-19 diagnosis. Lines between dots connect the individual HCW’s data points over the time course. Comparisons between groups in panels **(A and B)** were made using a two-way ANOVA with Bonferroni post-test. P-values are noted as **p<0.01, ***p<0.001, ****p<0.0001, and ns=p>0.05.

Additionally, we examined those HCWs that experienced breakthrough infection during Omicron subvariant waves in Columbus, Ohio. These Omicron-wave infected HCWs exhibited significantly increased nAb titers against BA.1, BA.2.12.1, and BA.4/5 at the 4-6 and 7-9 month timepoints (**Fig 3B**). Within our cohort, 13 HCWs provided samples in all three collection windows. Within this subset, those without any breakthrough infection largely exhibited declining nAb titers throughout the study period, while those who experienced breakthrough infection often recovered higher nAb titers (**Fig 3C-F**). Together, these results highlight the impact of breakthrough infection, particularly by Omicron subvariants, in enhancing the strength and breath of boosted vaccinees nAb response.

### Administration of a second booster recovers neutralizing antibody titers

Two HCWs in our cohort were administered a second booster dose of mRNA vaccine about 7 months after receiving their first booster dose (**Table S2**). These two HCWs exhibited higher nAb titers at 2-3 weeks post first booster vaccination but showed a strong decline in nAb titers against all variants tested at about 4 months after receiving the first booster (**Fig 4**). The decline was most dramatic against the Omicron subvariants compared to D614G, with NT_50_ values below the limit of detection (**Fig 4A, B, D, E, G**). Remarkably, the administration of a second booster dose was able to recover nAb titers to levels comparable with the ∼2-3 week-post 1^st^ booster timepoint, with a sample being taken approximately 2 weeks post-second booster (**Fig 4C, F, G**). Hence, second booster vaccination is needed to restore nAb levels in individuals, most of whom experience dramatic declines following the first booster.

**Figure 4:**
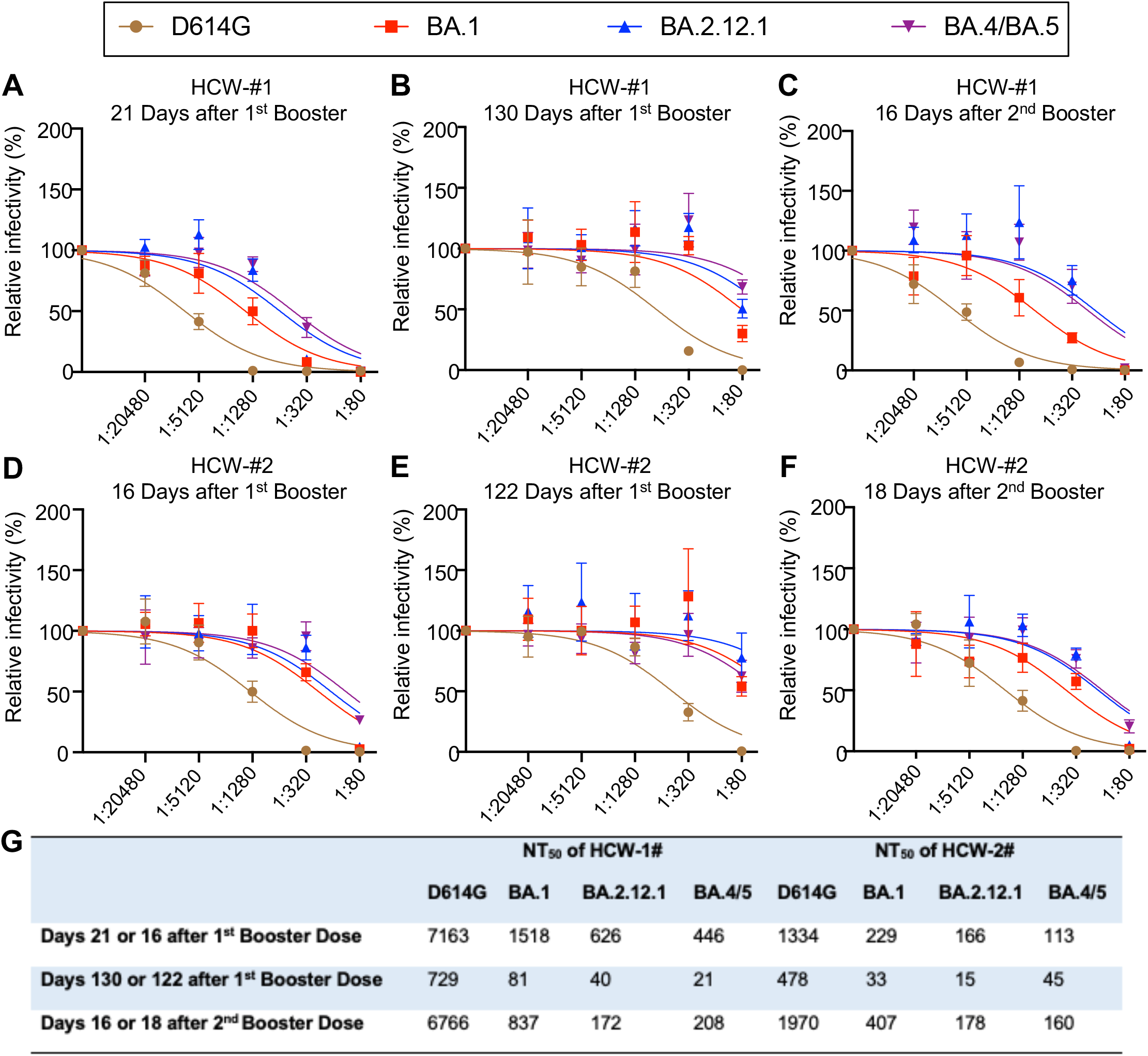
Administration of a second mRNA booster dose recovers nAb titers. Neutralization curves are depicted for two HCWs (designated as HCW #1 and HCW #2) in the cohort that received a second mRNA booster vaccination. Individual curves represent the different variants tested (D614G, Omicron subvariants BA.1, BA.2.12.1, and BA.4/5). Days after administration of the first booster dose are listed in panels **(A), (B), (D), and (E)** while days after administration of the second booster are listed in **(C and F)**. Both HCWs received the second booster about 7 months after the first booster dose. **(G)** Table summarizing the NT_50_ values for each HCW against each of the variants at timepoints pre- and post-second booster.

## Discussion

To maintain protection against severe outcomes of COVID-19, it is critical to understand how well and for how long booster vaccination induces robust levels of nAb against emerging variants of SARS-CoV-2. In this study, we determined the durability of mRNA booster vaccination in our cohort of HCWs against the most recent Omicron subvariants BA.2.12.1, and BA.4/5, along with original BA.1 and prototype D614G. In agreement with our previous reports, we demonstrate a marked reduction in nAb titer against Omicron subvariants BA.1, BA.2.12.1, and BA.4/5 relative to D614G for post booster vaccination samples (Evans et al., 2022c, 2022b; Qu et al., 2022).

Critically, our data demonstrates a modest decline, with ∼2.5-fold decrease at 7-9 month following booster vaccination and 10-14% of nAb titer drop every 30 days, in booster durability over time for all variants tested. This is in sharp contrast to the 7-10 fold decrease in nAb titer 6 months after second mRNA dose (Evans et al., 2022a), demonstrating a much more stable nAb titer than that provided by two doses of mRNA vaccine. These results are consistent with reports of a reduction in vaccine efficacy against infection and hospitalization up to four months post-booster dose (Ferdinands et al., 2022; Richterman et al., 2022). The rate of decay suggests that at least an annual booster vaccine may be required to provide sufficient protection against COVID-19 in the coming years. Notably, the rate of decline in booster durability appears largely consistent between each of the variants tested, including the prototype D614G and original Omicron BA.1. However, the Omicron subvariants exhibit strong neutralization resistance overall.

Breakthrough infections represent an additional antigenic exposure that can further boost nAb titers, especially against variants similar to the one with which the individual was infected. We observed that HCWs who tested positive for COVID-19 exhibit significant boosts in nAb titers for all variants tested and at nearly every time point. Importantly, this effect appears most robust against the Omicron subvariants (up to 8-fold) when compared to D614G (about 2-3 fold), especially for those HCWs that were infected during Omicron waves. These data suggest that an Omicron-specific antigenic exposure provides a critical boost to nAb titers against BA.1, BA.2.12.1, and BA.4/5.

Overall, we demonstrate that booster durability declines over time, but to a much lesser extent compared to the decay of nAb provided by two doses of mRNA vaccine alone. While the rate of booster nAb durability decay is similar among variants, the Omicron subvariants, especially BA.4/5, exhibit substantial neutralization resistance. This may suggest that SARS-CoV-2 variant evolution leading to immune evasion in a HCW cohort plays a more critical role in determining booster efficacy than the passage of time, given the modest waning of booster-induced immunity. It should be noted that the relative waning of booster induced immunity in vulnerable populations, including the elderly, remains to be investigated. Our study suggests a fourth dose of vaccine, or second booster, and perhaps an Omicron-specific booster, may become necessary. As new variants evolve, vaccine reformulation may be required to maintain sufficient protection from emerging strains.

## Acknowledgements

We thank the NIH AIDS Reagent Program and BEI Resources for providing important reagents for this work. We also thank the Clinical Research Center/Center for Clinical Research Management of The Ohio State University Wexner Medical Center and The Ohio State University College of Medicine in Columbus, Ohio, specifically Francesca Madiai, Dina McGowan, Breona Edwards, Evan Long, and Trina Wemlinger, for logistics, collection and processing of samples. In addition, we thank Sarah Karow, Madison So, Preston So, Daniela Farkas, and Finny Johns in the clinical trials team of The Ohio State University for sample collection and other supports. This work was supported by a fund provided by an anonymous donor to Ohio State University (to Dr. Liu); an award (U54CA260582, to Drs. Lozanski, Saif, Oltz, Gumina, and Liu) from the National Cancer Institute of the National Institutes of Health (NIH); a grant (R01 AI150473, to Dr. Liu) from the NIH; a Glenn Barber Fellowship (to Mr. Evans) from the Ohio State University College of Veterinary Medicine; grants (to Dr. Gumina) from the Robert J. Anthony Fund for Cardiovascular Research and the JB Cardiovascular Research Fund; and a grant (R01 HD095881, to Dr. Saif) from the NIH.

## Author Contributions

S.-L.L. conceived and directed the project. P.Q. performed most of the experiments. J.N.F, J.P.E. assisted in experiments and contributed data processing and analyses. C.C., G.L., R.J.G. provided clinical samples. P.Q., J.N.F., J.P.E., and S.-L.L. wrote the paper. Y.-M.Z, L.J.S., E.M.O., and R.J.G. provided insightful discussion and revision of the manuscript.

## STAR Methods

### RESOURCE AVAILABILITY

#### Lead contact

Further information and requests for resources and reagents should be directed to the lead contact, Dr. Shan-Lu Liu.

#### Materials availability

Plasmids generated in this study are available upon request made to the lead contact.

#### Data and code availability

- NT50 values and de-identified health care worker information will be deposited to the National Cancer Institute SeroNet Coordinating Center. Additionally, NT50 values and de-identified patient information reported in this paper will be shared by the lead contact upon request.
- This paper does not report original code.
- Any additional information required to reanalyze the data reported in this paper is available from the lead contact upon request.

### EXPERIMENTAL MODEL AND SUBJECT DETAILS

#### Vaccinated cohort

Summary data on the HCW cohort is available in supplementary Table S1 and Table S2. Vaccinated HCW samples were collected under approved IRB protocols (2020H0228 and 2020H0527). In the study group, 46 HCWs received homologous vaccine and booster doses. Sera were collected at 3-month intervals after receiving the second dose of mRNA vaccine. Booster dose administration was variable within the study period resulting in sample collections occurring 1-9 months post booster dose administration, which are divided into 3 groups, i.e., 1-3 month, 4-6 month and 7-9 month with a total of 101 post-booster samples. These samples included 24 Moderna mRNA-1273 and 22 Pfizer/BioNTech BNT162b2 boosted HCWs. Dates of prior COVID-19 diageneses were self-reported.

#### Cell lines and maintenance

Human embryonic kidney cell line HEK293T (ATCC CRL-11268, RRID: CVCL_1926) and HEK293T overexpressing human ACE2 (BEI NR-52511, RRID: CVCL_A7UK) were cultured in DMEM (Gibco, 11965-092) supplemented with 10% FBS (Sigma, F1051) and 0.5% penicillin-streptomycin (HyClone, SV30010). Cells were maintained in 10cm dishes and incubated at 37°C and 5.0% CO_2_. For passaging, cells were first washed in Dulbecco’s phosphate buffer saline (Sigma, D5652-10X1L), and incubated in 0.05% Trypsin + 0.53 mM EDTA (Corning, 25-052-CI) for detachment.

## METHOD DETAILS

### Plasmids

Our pseudotyped lentiviral stocks were produced using our previously reported HIV-1-based vector (pNL4-3-inGluc) carrying a *Gaussia* luciferase reporter gene that is expressed and secreted by virally infected cells(Goerke et al., 2008; Mazurov et al., 2010; Zeng et al., 2020). SARS-CoV-2 spike constructs were generated and cloned into the pcDNA3.1 plasmid backbone using KpnI and BamHI restriction enzyme cloning by GenScript Biotech (Piscataway, NJ). These spike constructs bear N- and C-terminal FLAG tags.

### Pseudotyped lentivirus production

Pseudotyped lentiviral vectors were produced as previously reported (Evans et al., 2022a). HEK293T cells were transfected with the pNL4-3-inGluc vector alongside the spike construct of interest in a 2:1 ratio using polyethyleneimine transfection. Virus produced by the cells was harvested by collecting and replacing the culture media 48-, and 72-hours post-transfection. The relative infectivity of the viruses was determined in HEK293T-ACE2 cells by measuring *Gaussia* luciferase activity 48- and 72-hours post-infection; equivalent infectious viruses for each variant were used for the neutralization assay. Luciferase assays were conducted by taking a 20μL sample of infected cell culture media and combining it with 20μL of *Gaussia* luciferase substrate (0.1 M Tris pH 7.4, 0.3 M sodium ascorbate, 10 μM coelenterazine) and immediately measuring luminescence using a BioTek Cytation5 plate reader with Gen5 Microplate Reader and Imager Software version 3.03.

### Virus neutralization assay

Neutralization assays using pseudotyped lentiviral vectors were conducted as previously described (Evans et al., 2022a; Zeng et al., 2020, 2021b, 2021a) HCW serum samples were first serially diluted 4-fold (final dilutions 1: 80, 1:320, 1:1280, 1:5120, 1:20480, and no serum control) and combined with equal amounts of SARS-CoV-2 pseudotyped vector. The diluted sera and vector mix was then incubated 1 hour at 37°C then used to infect HEK293T-ACE2 cells. *Gaussia* luciferase activity was assessed 48- and 72-hours post infection as described in the previous section. NT50 values were determined by least-squares-fit, non-linear regression in GraphPad Prism 9 (San Diego, CA).

## QUANTIFICATION AND STATISTICAL ANALYSIS

All statistical analysis was performed using GraphPad Prism 9 and are described in the figure legends. NT_50_ values were determined by least-squares fit non-linear regression in GraphPad Prism 9. Throughout, statistical significance was determined using log_10_ transformed NT_50_ values to better approximate normality. Bars represent geometric means with 95% confidence intervals (Fig. 1A-C) and indicate means ± SEM (Fig 3A-B, S2). Generally, comparisons between multiple groups were made using a one-way ANOVA with Bonferroni post-test (Fig. 1A-C, S1) or two-way ANOVA with Bonferroni post-test (Fig. 3 A-B). Correlative analysis of nAb titers and time post booster dose administration was made using a least-squares fit linear regression model (Fig. 2A-D). Comparisons between two-groups were made using a two-tailed student’s t-test with Welch’s correction (Fig. S2 A-B). Due to small sample sizes, analysis of the influence of sex could not be performed without the influence of confounding variables including vaccination status, vaccine type, and time since vaccination skewing the analysis.

---

**Table S1:** Demographic and sample collection information of HCWs

|  | 1~3 months | 4~6 months | 7~9 months |
| --- | --- | --- | --- |
| **Age in Years at Sample Collection**<br>[Median (Range)] | 35 (26-61) | 35 (25-58) | 36.5 (25-61) |
| **Gender** [n (% of Total)] |  |  |  |
| Male | 20 (71.4%) | 20 (54.1%) | 21 (58.3%) |
| Female | 8 (28.6%) | 17 (45.9%) | 15 (41.7%) |
| **Sample Collection Window** | Oct. 2021-Feb. 2022 | Feb. 2022-Jun. 2022 | Feb. 2022-Jun. 2022 |
| **Type of 1<sup>st</sup> Booster Dose Vaccine** [n (% of Total)] |  |  |  |
| Pfizer | 17 (60.7%) | 17 (45.9%) | 17 (47.2%) |
| Moderna | 11 (39.3%) | 20 (54.1%) | 19 (52.8%) |
| **Sample Collection Timing** [Median (Range)] |  |  |  |
| Days post 1<sup>st</sup> Booster Dose | 42 (7-86) | 112 (92-178) | 212.5 (182-283) |
| **Prior SARS-CoV-2 Infection Confirmed by PCR** [n (% of Total)] |  |  |  |
| Prior Omicron Wave Infection | 4 (14.3%) | 4 (10.8%) | 3 (8.3%) |
| Omicron Wave Infection | 2 (7.1%) | 5 (13.5%) | 8 (22.2%) |

Summary information for the HCW sera samples collected post 1<sup>st</sup> booster dose of mRNA vaccine is shown.

**Table S2:** Sample collection information of two HCWs receiving 2^nd^ booster dose of vaccine

|  | HCW-#1 | HCW-#2 |
| --- | --- | --- |
| **Gender** | Female | Male |
| **Age at Collection** | 48 | 48 |
| **COVID-19 Status** | Negative | Negative |
| **Type of Vaccine** |  |  |
| Type of 1<sup>st</sup> Booster Dose Vaccine | Pfizer | Pfizer |
| Type of 2<sup>nd</sup> Booster Dose Vaccine | Pfizer | Pfizer |
| **Sample Collection Timing** |  |  |
| Days Post 1<sup>st</sup> booster dose (Days Pre 2<sup>nd</sup> booster dose) | 21 (215) | 16 (194) |
| Days Post 1<sup>st</sup> booster dose (Days Pre 2<sup>nd</sup> booster dose) | 130 (106) | 122 (88) |
| Days Post 1<sup>st</sup> booster dose (Days Post 2<sup>nd</sup> booster dose) | 252 (16) | 228 (18) |

Summary information of the sera samples of two HCWs who received two booster doses of vaccine is shown.

**Figure S1:**
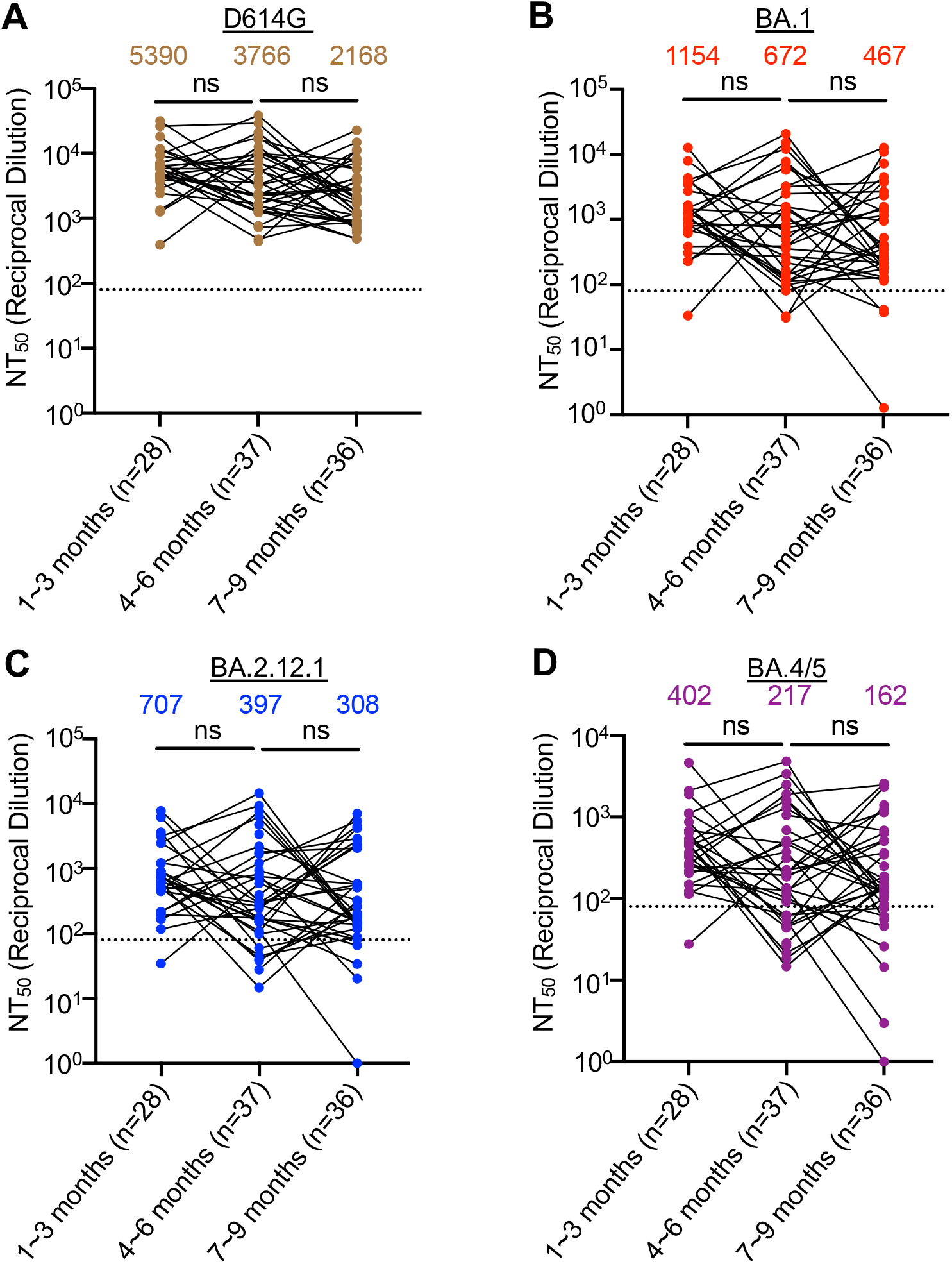
Booster vaccination-induced nAb response exhibits modest waning over time, related to Figure 1. Here, nAb titers in serum samples from HCWs collected at 1-3, 4-6, and 7-9 months after receiving a booster dose of mRNA vaccine are sorted by on the variant being tested for **(A)** D614G, **(B)** BA.1, **(C)** BA.2.12.1, and **(D)** BA.4/5. Dots represent individual samples; lines connect dots that were from the same individual HCW. Significance values were determined using a one-way ANOVA. P-values are represented as ns=not significant.

**Figure S2:**
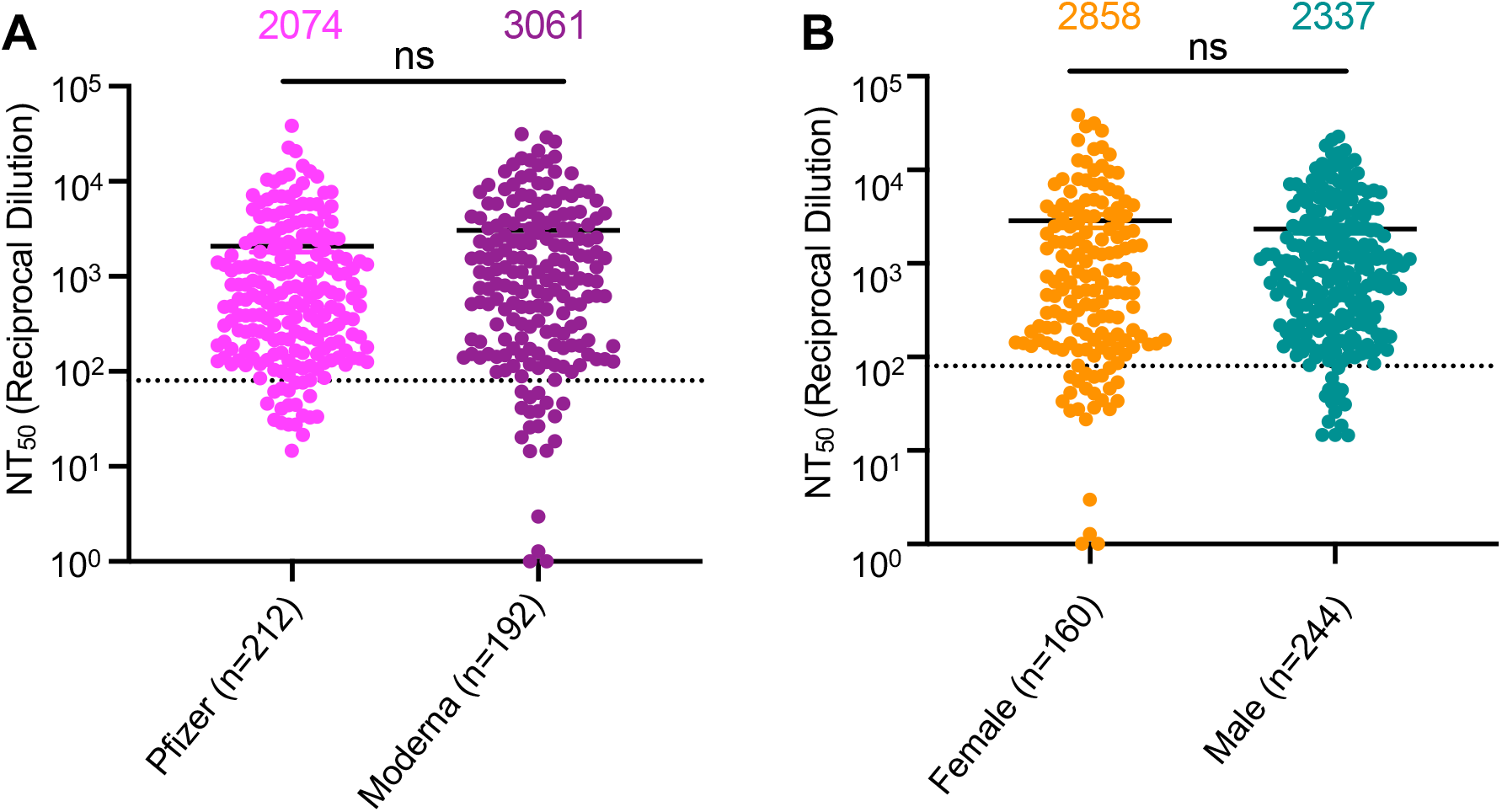
Booster vaccination-induced nAb response does not vary between vaccine manufacturer or HCW sex, related to Figure 1. Displayed are the nAb titers for sera from HCWs for all 3 timepoints (1-3 months, 4-6 months, and 7-9 months) and all 4 variants (D614G, BA.1, BA.2.12.1, and BA.4/5) pooled together and separated by **(A)** vaccine manufacturer and **(B)** sex. Comparisons were made using a two-tailed student’s t-test with Welch’s correction. P-values are represented as ns=not significant.

## References

Andrews, N., Stowe, J., Kirsebom, F., Toffa, S., Sachdeva, R., Gower, C., Ramsay, M., and Lopez Bernal, J. (2022). Effectiveness of COVID-19 booster vaccines against COVID-19-related symptoms, hospitalization and death in England. Nature Medicine 28, 831–837. https://doi.org/10.1038/s41591-022-01699-1.

Centers for Disease Control and Prevention (2022). Variant Proportions.

Chenchula, S., Karunakaran, P., Sharma, S., and Chavan, M. (2022). Current evidence on efficacy of COVID-19 booster dose vaccination against the Omicron variant: A systematic review. Journal of Medical Virology 2969–2976. https://doi.org/10.1002/jmv.27697.

Evans, J.P., Zeng, C., Carlin, C., Lozanski, G., Saif, L.J., Oltz, E.M., Gumina, R.J., and Liu, S.L. (2022a). Neutralizing antibody responses elicited by SARS-CoV-2 mRNA vaccination wane over time and are boosted by breakthrough infection. Science Translational Medicine 14. https://doi.org/10.1126/scitranslmed.abn8057.

Evans, J.P., Qu, P., Zeng, C., Zheng, Y.-M., Carlin, C., Bednash, J.P., Lozanski, G., Mallampalli, R., Saif, L.J., Oltz, E.M., et al. (2022b). Neutralization of the SARS-CoV-2 Deltacron and BA.3 Variants. New England Journal of Medicine.

Evans, J.P., Zeng, C., Qu, P., Faraone, J., Zheng, Y.-M., Carlin, C., Bednash, J.S., Zhou, T., Lozanski, G., Mallampalli, R., et al. (2022c). Neutralization of SARS-CoV-2 Omicron Sub-lineages BA.1, BA.1.1, and BA.2. Cell Host & Microbe https://doi.org/10.1016/j.chom.2022.04.014.

Ferdinands, J.M., Rao, S., Dixon, B.E., Mitchell, P.K., DeSilva, M.B., Irving, S.A., Lewis, N., Natarajan, K., Stenehjem, E., Grannis, S.J., et al. (2022). Waning 2-Dose and 3-Dose Effectiveness of mRNA Vaccines Against COVID-19–Associated Emergency Department and Urgent Care Encounters and Hospitalizations Among Adults During Periods of Delta and Omicron Variant Predominance — VISION Network, 10 States, Aug. MMWR Recommendations and Reports 71, 255–263. https://doi.org/10.15585/mmwr.mm7107e2.

Ghimire, D., Han, Y., and Lu, M. (2022). Structural Plasticity and Immune Evasion of SARS-CoV-2 Spike Variants. Viruses 14, 1255. https://doi.org/10.3390/v14061255.

Goerke, A.R., Loening, A.M., Gambhir, S.S., and Swartz, J.R. (2008). Cell-free metabolic engineering promotes high-level production of bioactive Gaussia princeps luciferase. Metabolic Engineering 10, 187–200. https://doi.org/10.1016/j.ymben.2008.04.001.

Mazurov, D., Ilinskaya, A., Heidecker, G., Lloyd, P., and Derse, D. (2010). Quantitative comparison of HTLV-1 and HIV-1 cell-to-cell infection with new replication dependent vectors. PLoS Pathogens 6. https://doi.org/10.1371/journal.ppat.1000788.

O’Toole, Á., Scher, E., Underwood, A., Jackson, B., Hill, V., McCrone, J.T., Colquhoun, R., Ruis, C., Abu-Dahab, K., Taylor, B., et al. (2021). Assignment of epidemiological lineages in an emerging pandemic using the pangolin tool. Virus Evolution 7, 1–9. https://doi.org/10.1093/ve/veab064.

Qu, P., Faraone, J., Evans, J.P., Zou, X., Zheng, Y.-M., Carlin, C., Bednash, J.S., Lozanski, G., and Mallampalli, R.K. (2022). Neutralization of the SARS-CoV-2 Omicron BA.4/5 and BA.2.12.1 Subvariants. New England Journal of Medicine https://doi.org/10.1056/NEJMc2206725.

Rajpal, V.R., Sharma, S., Sehgal, D., Singh, A., Kumar, A., Vaishnavi, S., Tiwari, M., Bhalla, H., Goel, S., and Raina, S.N. (2022). A comprehensive account of SARS-CoV-2 genome structure, incurred mutations, lineages and COVID-19 vaccination program. Future Virology https://doi.org/10.2217/fvl-2021-0277.

Richterman, A., Behrman, A., Brennan, P.J., O’Donnel, J.A., Snider, C.K., and Chaiyachati, K.H. (2022). Durability of SARS-CoV-2 mRNA Booster Vaccine Protection Against Omicron Among Health Care Workers with a Vaccine Mandate. Clinical Infectious Diseases https://doi.org/10.1093/cid/ciac454.

Scobie, H.M., Johnson, A.G., Suthar, A.B., Severson, R., Alden, N.B., and Balter, S. (2021). Monitoring Incidence of COVID-19 Cases, Hospitalizations, and Deaths, by Vaccination Status — 13 U.S. Jurisdictions, April 4–July 17, 2021. CDC Morbidity and Mortality Weekly Report 70, 1284–1290..

WHO (2021). WHO Coronavirus (COVID-19) Dashboard.

Zeng, C., Evans, J.P., Pearson, R., Qu, P., Zheng, Y.M., Robinson, R.T., Hall-Stoodley, L., Yount, J., Pannu, S., Mallampalli, R.K., et al. (2020). Neutralizing antibody against SARS-CoV-2 spike in COVID-19 patients, health care workers, and convalescent plasma donors. JCI Insight 5. https://doi.org/10.1172/jci.insight.143213.

Zeng, C., Evans, J.P., Faraone, J.N., Qu, P., Zheng, Y.M., Saif, L., Oltz, E.M., Lozanski, G., Gumina, R.J., and Liu, S.L. (2021a). Neutralization of SARS-CoV-2 Variants of Concern Harboring Q677H. MBio 12. https://doi.org/10.1128/mBio.02510-21.

Zeng, C., Evans, J.P., Reisinger, S., Woyach, J., Liscynesky, C., Boghdadly, Z. el, Rubinstein, M.P., Chakravarthy, K., Saif, L., Oltz, E.M., et al. (2021b). Impaired neutralizing antibody response to COVID-19 mRNA vaccines in cancer patients. Cell and Bioscience 11. https://doi.org/10.1186/s13578-021-00713-2.

